# Conservation of Long G4-rich (LG4) genomic enhancer regulations

**DOI:** 10.64898/2026.03.11.711068

**Authors:** Michael H. Shaw, Jeffrey D. DeMeis, Claire A. Arnold, Makala R. Cox, Tien C. Duong, Kyle A. Gaviria, Grace K. McDavid, Jose M. Villegas, Margaret L. Weimer, Suhas S. Patil, Shahem Y. Alqudah, Glen M. Borchert

## Abstract

Long G4-rich regions (LG4s) are defined as DNA sequences containing a high density of guanine triplets capable of forming non-B DNA structures called G-quadruplexes (G4s). These regions frequently overlap with enhancers, which are regulatory DNA elements that modulate gene expression by interacting with DNA regions that dictate where transcription is initiated known as promoters. While LG4s have now been well-characterized in the human genome, neither LG4 occurrence, nor the ability of LG4s to function as enhancers, in other species has been described. To address this, we screened the genomes of 16 different species from various taxa to identify LG4s and then determined if they were conserved, and if so, if their regulatory capacity was similarly conserved. Our analyses characterized a number of previously unreported LG4s in the human genome as well as LG4s in 13 additional species. Of note, we identified a highly conserved LG4 enhancer predicted to regulate over 40 genes. This LG4 is embedded in the *MAZ* (Myc-Associated Zinc finger protein) locus, and we find this LG4 possesses the ability to directly interact with the same target promoter in both human and mouse. In summary, this work describes LG4s in the genomes of both unicellular and multicellular species including vertebrates, invertebrates, plants, and fungi. Furthermore, many of these LG4 sequences are highly conserved as is their regulatory capacity.

## Introduction

G-quadruplexes (G4s) are non-B form, single stranded DNA structures formed by hydrogen bonding between guanine (G) nucleotides (**Figure 1A**) and are minimally defined by the sequence motif GGGnGGGnGGGnGGG where ‘n’ corresponds to a spacer sequence of variable length and nucleotide composition^1-4^ (**Figure 1B**). These G4 motifs are ubiquitously found in the human genome and are even reported to be present in more primitive taxonomies including viridae^5^. In 2020, we identified 301 long genomic stretches significantly enriched for minimal G4 motifs (LG4s) in humans. Importantly, many of these LG4s were determined to overlap human regulatory elements such as gene promoters and enhancers. In fact, 217/301 of these LG4s overlap an annotated human enhancer element^6^.

**Figure 1.**
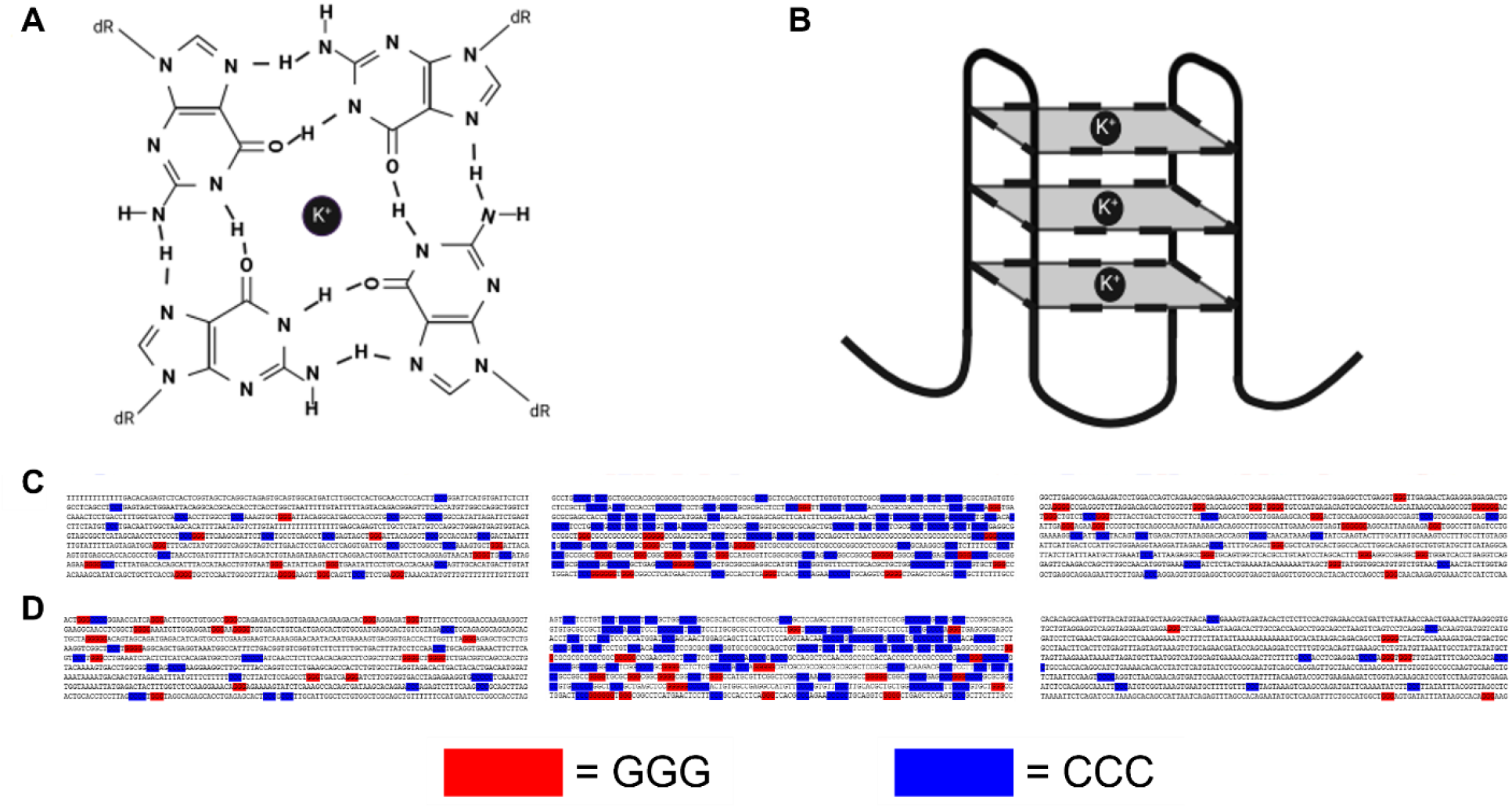
LG4 DNA located within *MAZ* loci in human and mouse. (**A**) Structure of an individual G-quartet wherein guanine nucleotides are held together by hydrogen bonds centered about a central potassium cation (K^+^). (**B**) Cartoon depiction of unimolecular antiparallel G4 DNA where each square corresponds to an individual G-quartet and the corners of each quartet correspond to a single guanine. (**C**) Human *MAZ* LG4 locus in which ≥3 consecutive genomic Gs (guanines) and ≥3 consecutive genomic Cs (cytosines) are highlighted. Top center, LG4 located at human Chr16:29805901:29806863; Top left and right, ~1 kb sequences located 40 kb upstream and downstream of the central LG4 (Chr16:29765901:29766863 and Chr16:29845901:29846863, respectively). (**D**) Mouse *MAZ* LG4 locus in which ≥3 consecutive genomic Gs and ≥3 consecutive genomic Cs are highlighted. Bottom center, LG4 located at mouse Chr7:126626008:126626970. Bottom left and right, ~1 kb sequences located 40 kb upstream and downstream of the central LG4 (Chr7:126585701:126586663 and Chr7:126665701:126666663, respectively). **A** and **B** adapted from DeMeis et al., 2025.

Gene enhancers constitute functional regulatory elements capable of modulating the activities of distally and/or proximally located cognate gene promoters to regulate target gene expression. Historically, the interaction between gene enhancers and their respective target promoters were believed to proceed through chromatin looping facilitated by protein interactions occurring between separate protein complexes bound to the enhancer and promoter elements^7-9^. That said, we recently found that an LG4 enhancer on human Chr5 and an LG4 enhancer on human Chr12 associate with their target promoters through direct DNA::DNA interaction between G4 sequences embedded within those regulatory elements^10^.

The existence of LG4s in other species, however, has yet to be determined. Given that functionally relevant genetic regulatory elements, such as gene promoters are often evolutionarily conserved^11^, and the significant association of LG4s with human regulatory regions, we postulated that LG4s are broadly conserved across species. As such, this work describes our screening of a number of genomes for LG4s, comparative analyses examining LG4 conservation, and biochemical assessment of the ability of select, conserved LG4 enhancers (**Figure 1C, D**) to directly interact with target promoters in both humans and mice.

## Methods

### LG4 identification

The genomes of 16 different species were analyzed: *Homo sapiens* (GRCh38.p14), *Xenopus tropicalis* (UCB_Xtro_10.0), *Zea mays* (Zm-B73-REFERENCE-NAM-5.0), *Macaca mulatta* (Mmul_10), *Gallus gallus* (bGalGal1.mat.broiler.GRCg7b), *Mus musculus* (GRCm39), *Chlamydomonas reinhardtii* (Chlamydomonas_reinhardtii_v5.5), *Sus scrofa* (Sscrofa11.1), *Danio rerio* (GRCz11), *Drosophila melanogaster* (BDGP6.46), *Caenorhabditis elegans* (WBcel235), *Schizophyllum commune* (GCA000143185v1), *Neurospora crassa* (NC12), *Arabidopsis thaliana* (TAIR10), *Haloferax volcanii* (ASM2568v1), and *Saccharomyces cerevisiae* (R64-1-1). LG4ID was performed in Python and is available at GitHub^12^: https://github.com/glen-borchert/LG4ID. LG4s were minimally defined as 500 bp sequences possessing at least 40 occurrences of ‘GGG’ or 40 occurrences of ‘CCC’. LG4ID sectioned genomes into 10 Mb bins then analyzed these via a 500 bp sliding window for sequences meeting the LG4 inclusion criteria. When a sequence meeting the G/C triplet threshold was identified, the program extracted the 500 bp sequence and extended the 3’ end until the minimum G/C triplet density fell below 40 G or C triplets/500 bp at which point the program deposited the full length LG4 into a final output file. If the entire 3’ flanking sequence met the inclusion criteria, then the program would continue to examine and include sequence from the subsequent bin. To ensure the LG4s identified were not telomere repeats, any LG4 possessing ≥4 iterations of the telomere repeat sequence corresponding to that species was removed from consideration.

### LG4 informatic analysis

OverlapID was performed in Python (available at: https://github.com/glen-borchert/OverlapID) in order to determine overlaps between LG4 genomic positions and the positions of annotated enhancers and genes. Gene positions for all species were obtained via Ensembl BioMart^13^, except for *Schizophyllum commune*, which was obtained via Ensembl^14^ FTP. All available promotor and enhancer positions from EnhancerAtlas^15^, Ensembl BioMart^13^, and GeneHancer^16^ were collected for each species (**Table 1**). Evolutionary conservation of LG4s was assessed by performing sequence alignments using BLAST (ncbi-blast-2.16.0+-win64)^17^ with sequences possessing >80% identity across at least 50 bp considered conserved.

**Table 1.** LG4s identified in select species. LG4 #, the number of LG4s identified via LG4ID; Genome Size (bp), genome length in current Ensembl^14^ assembly; (LG4)PM, LG4# Per Million bp; Genes, number of overlapping genes identified via OverlapID; Promoters, number of overlapping promoters identified via OverlapID; Enhancers, number of overlapping enhancers identified via OverlapID. “d/u” indicates data unavailable.

| Species Name | LG4 # | Genome Size (bp) | (LG4)PM | Genes | Promoters | Enhancers |
| --- | --- | --- | --- | --- | --- | --- |
| *Xenopus tropicalis* | 10,507 | 1,451,301,209 | 7.24 | 404 | d/u | d/u |
| *Zea mays* | 9,574 | 2,182,075,994 | 4.39 | 39 | d/u | d/u |
| *Macaca mulatta* | 2,753 | 2,971,331,530 | 0.93 | 223 | d/u | d/u |
| *Gallus gallus* | 2,523 | 1,053,332,251 | 2.40 | 528 | 164 | 373 |
| *Homo sapiens* | 2,278 | 3,099,750,718 | 0.73 | 272 | 366 | 1,076 |
| *Mus musculus* | 2,015 | 2,728,222,451 | 0.74 | 46 | 34 | 203 |
| *Chlamydomonas reinhardtii* | 917 | 111,098,438 | 8.25 | 24 | d/u | d/u |
| *Sus scrofa* | 648 | 2,501,912,388 | 0.26 | 89 | 233 | 68 |
| *Danio rerio* | 154 | 1,373,471,384 | 0.11 | 0 | d/u | 87 |
| *Drosophila melanogaster* | 11 | 143,726,002 | 0.08 | 0 | d/u | 0 |
| *Caenorhabditis elegans* | 8 | 100,286,401 | 0.08 | 2 | 0 | 3 |
| *Schizophyllum commune* | 8 | 38,670,379 | 0.21 | 0 | d/u | d/u |
| *Neurospora crassa* | 1 | 41,102,378 | 0.02 | 0 | d/u | d/u |
| *Arabidopsis thaliana* | 0 | 119,667,750 | 0.00 | d/u | d/u | d/u |
| *Haloferax volcanii* | 0 | 4,012,900 | 0.00 | d/u | d/u | d/u |
| *Saccharomyces cerevisiae* | 0 | 12,157,105 | 0.00 | d/u | d/u | d/u |

### Plasmid construction

The human *KIF22* promoter and human *MAZ* LG4 were each PCR amplified using 10 ng of WI2-3752P18 fosmid DNA as template, whereas the mouse *Kif22* promoter and mouse *MAZ* LG4 were each amplified using 10 ng of genomic mouse DNA as template. WI2-3752P18 fosmid (BACPAC Genomics, Inc. Redmond, WA) was isolated by HighPrep Plasmid DNA Kit (Magbio Genomics Inc. Gaithersburg, MD 501656596). PCRs were performed using LongAmp Taq DNA polymerase (NEB 50994936) in 25 μL according to manufacturer’s protocol with 5% DMSO. Notably, nested PCRs were performed to clone all promoters and LG4s in human and mouse. For each construct amplified by nested PCR, the primers listed in **Supplementary Table 5** are numbered to indicate reaction order. Resulting amplicons were gel extracted using Wizard SV Gel and PCR Clean-Up System (Promega PR-A9281) then cloned into TOPO TA pcR2.1 (Invitrogen 45–064-1) and verified by sequencing (Eurofins USA Lancaster, PA).

### Electrophoretic Mobility Shift Assays

LG4 and promoter sequences used in electrophoretic mobility shift assays (EMSAs) were cloned into TOPO TA pCR2.1 (Invitrogen 45–064-1) in both orientations and confirmed by sequencing. Confirmed constructs were transformed into *F’Iq* Competent E. coli (NEB C2992H) and colonies picked to inoculate 50 mL of LB then grown at 37°C for 4 hr at 225 RPM in an orbital shaker. M13KO7 helper phage (NEB N0315S) was then added to each [1 × 10^8^ pfu/ml] and incubated at 37°C for 1.5 hr at 225 RPM then kanamycin added [70 μg/mL] and cultures allowed to grow overnight at 37°C at 225 RPM. Single-stranded DNA (ssDNA) was isolated 18 hr later per M13KO7 helper phage manufacturer protocol (NEB N0315S).

ssDNA constructs were then run at 90 V for 1.25 hr on a on a 1% agarose 1× TBE gel then ssDNA size selected and purified using Wizard SV Gel and PCR Clean-Up System (Promega, Madison, WI, PR-A9281) then quantified using a Nanodrop 6000 (Thermo-Scientific). A total of 20 μL [20 ng/μL] of each ssDNA promoter was boiled either separately or together with 10 μL [20 ng/μL] of LG4 ssDNA at 98°C for 10 min then held at 80°C for 10 min during which pre-heated KCl solution was added to bring the sample to 1 M final concentration (or an equal volume of 80°C ddH_2_O as control). Samples were then cooled to 45°C over 1 hr, then further cooled to 16°C over 1 additional hr, and then held at 4°C until use. Resulting ssDNA incubations were run on a 1.5% agarose 1× Tris-glycine (BioRad Hercules, CA, 1610734) gel at 75 V for 4 hr at 4°C, then gels were stained for 24 hr with SYBR Gold (Invitrogen S11494) diluted 1:10000 in 1×Tris-glycine (Bio-Rad 1610734). Finally, gels were imaged on a UV Transilluminator FBTIV-88 (Fisher Scientific).

## Results

### Identification of LG4s in select taxa

Using LG4ID, we identified 2,278 LG4s in the human genome (*Homo sapiens*, GRCh38) and find them to be relatively evenly distributed across chromosomes (excluding the majority of Chromosome Y and relatively large sections of Chromosomes 1, 3, 5, 12, and X all of which are largely devoid of LG4s.) (**Figure 2**). Of these human LG4s, 272 overlap annotated genes, 366 overlap promoters, and 1,076 overlap enhancers (Ensembl BioMart^13^) (**Table 1**) with 211 LG4s occupying genomic regions annotated as both promoters and enhancers (GeneHancer^16^).

**Figure 2.**
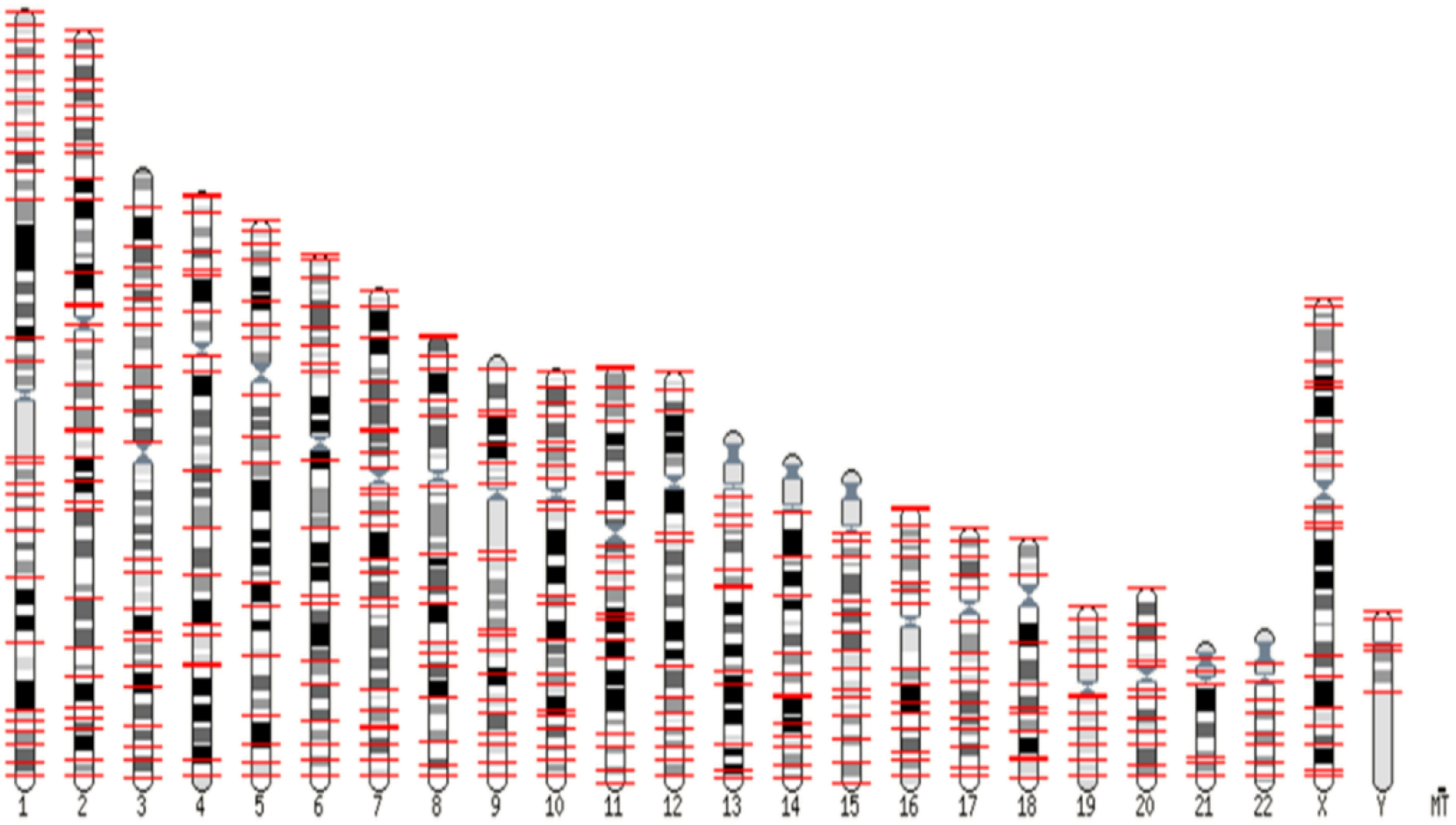
LG4 distribution across human chromosomes. Karyotypic view of all 2,278 LG4s identified within the human genome (GRCh38) showing their distribution across chromosomes. Red tick marks correspond to the approximate genomic location of each LG4. Image generated by Ensembl^14^.

As summarized in **Table 1**, we identified 10,507 LG4s in the *Xenopus tropicalis* genome (UCB_Xtro_10.0). While there are no currently available regulatory feature databases containing promoter and enhancer annotations for this species, 404 LG4s do overlap Ensembl BioMart^13^ annotated genes. In *Zea mays* (Zm-B73-REFERENCE-NAM-5.0), we identified 9,574 LG4s with 39 of these overlapping annotated genes (Ensembl BioMart^13^). In *Macaca mulatta* (Mmul_10), we identified 2,753 LG4s with 223 of these overlapping annotated genes (Ensembl BioMart^13^). In *Gallus gallus* (bGalGal1.mat.broiler.GRCg7b), we identified 2,523 LG4s with 528 overlapping annotated genes (Ensembl BioMart^13^), 164 overlapping annotated promoters (EnhancerAtlas^15^), and 373 overlapping annotated enhancers (EnhancerAtlas^15^). In *Mus musculus* (GRCm39), we identified 2,015 LG4s with 46 overlapping annotated genes, 34 overlapping annotated promoters, and 203 overlapping annotated enhancers (Ensembl BioMart^13^). In *Chlamydomonas reinhardtii* (Chlamydomonas_reinhardtii_v5.5), we identified 917 LG4s with 24 overlapping annotated genes (Ensembl BioMart^13^). In *Sus scrofa* (Sscrofa11.1), we identified 648 LG4s with 89 overlapping annotated genes (Ensembl BioMart^13^), 233 overlapping annotated promoters (EnhancerAtlas^15^), and 68 overlapping annotated enhancers (EnhancerAtlas^15^). In *Danio rerio* (GRCz11), we identified 154 LG4s with none of these overlapping annotated genes (Ensembl BioMart^13^), and 87 overlapping annotated enhancers (EnhancerAtlas^15^). In *Drosophila melanogaster* (BDGP6.46), we identified 11 LG4s, none of which were found to overlap annotated genes (Ensembl BioMart^13^) or regulatory features (EnhancerAtlas^15^). In *Caenorhabditis elegans* (WBcel235), we identified 8 LG4s with 2 overlapping annotated genes (Ensembl BioMart^13^), and 3 overlapping annotated promoters or enhancers (EnhancerAtlas^15^). In *Schizophyllum commune* (GCA000143185v1), we identified 8 LG4s with none of these overlapping annotated genes (Ensembl BioMart^13^). In *Neurospora crassa* (NC12), we identified only 1 LG4, and it did not overlap any annotated genes (Ensembl BioMart^13^). Interestingly, no LG4s were identified in the genomes of *Arabidopsis thaliana* (TAIR10), *Haloferax volcanii* (ASM2568v1), or *Saccharomyces cerevisiae* (R64-1-1). The list of all LG4s, their positions, and their sequences identified for each species can be found in **Supplemental Table 1**. The list of all enhancer and gene overlaps identified for each species can be found in **Supplemental Tables 2** and **3**, respectively.

### LG4s are conserved and associated with annotated regulatory regions and genes

Although we find no LG4s conserved across all taxa examined, we find 692 human LG4s to be conserved in at least one mammal, 26 in at least two, and six human LG4s conserved across all mammals analyzed (85.2%/793 bp). The full list of conserved LG4s can be found in **Supplemental Table 4**. Beyond conservation of LG4 sequences, we find several LG4 overlaps with annotated regulatory regions and/or genes are also conserved across species (**Supplemental Figures 1**-**7**). Particularly of note, we find conservation of the regulatory regions associated with the six LG4s common to human, mouse, pig, and rhesus monkey (**Table 2, Supplemental Figure 1**).

**Table 2.**
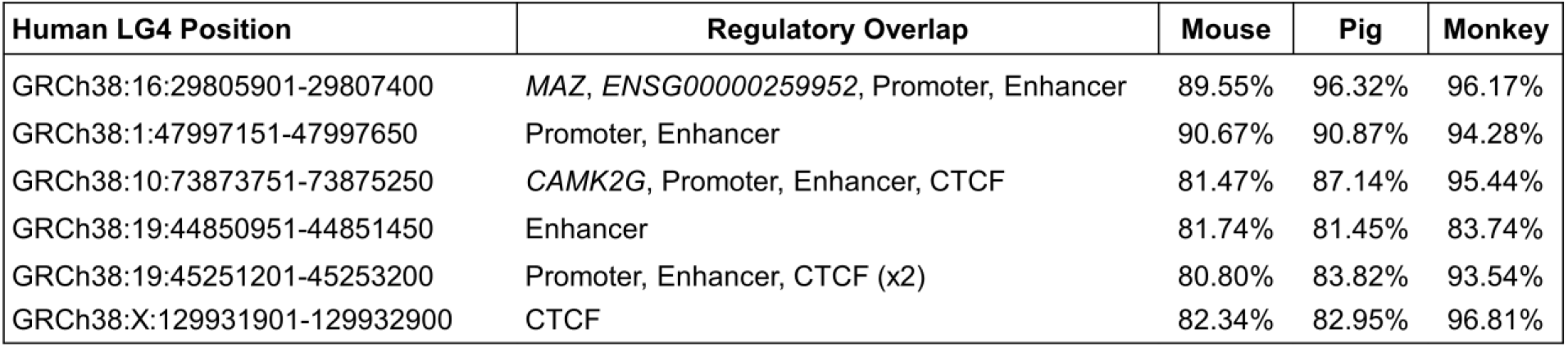
Conservation of the regulatory regions and/or genes associated with six LG4s common to human, mouse, pig, and rhesus monkey. Select conserved human LG4: regulatory element and/or gene overlaps. Percent identity of the human LG4 with each species is indicated. CTCF refers to CCCTC-binding Factor.

| Human LG4 Position | Regulatory Overlap | Mouse | Pig | Monkey |
| --- | --- | --- | --- | --- |
| GRCh38:16:29805901-29807400 | MAZ, ENSG00000259952, Promoter, Enhancer | 89.55% | 96.32% | 96.17% |
| GRCh38:1:47997151-47997650 | Promoter, Enhancer | 90.67% | 90.87% | 94.28% |
| GRCh38:10:73873751-73875250 | CAMK2G, Promoter, Enhancer, CTCF | 81.47% | 87.14% | 95.44% |
| GRCh38:19:44850951-44851450 | Enhancer | 81.74% | 81.45% | 83.74% |
| GRCh38:19:45251201-45253200 | Promoter, Enhancer, CTCF (x2) | 80.80% | 83.82% | 93.54% |
| GRCh38:X:129931901-129932900 | CTCF | 82.34% | 82.95% | 96.81% |

### *CAMK2G* LG4 is conserved across Mammalia

As a specific example of the LG4s listed in **Table 2**, an LG4 identified at human GRCh38:10:73873751–73875250 is conserved in the genomes of mouse, pig, and rhesus monkey (**Figure 3A-D**). In the human genome, this LG4 overlaps a known enhancer, promoter, CTCF site, and the *CAMK2G* (Calcium/Calmodulin-Dependent Protein Kinase II Gamma) gene. Notably, this LG4 similarly overlaps *CAMK2G* in mouse, pig, and rhesus monkey, and is associated with an enhancer and an EMAR (Epigenetically Modified Accessible Region)^14^ in both human and mouse (**Figure 3E, F**).

**Figure 3.**
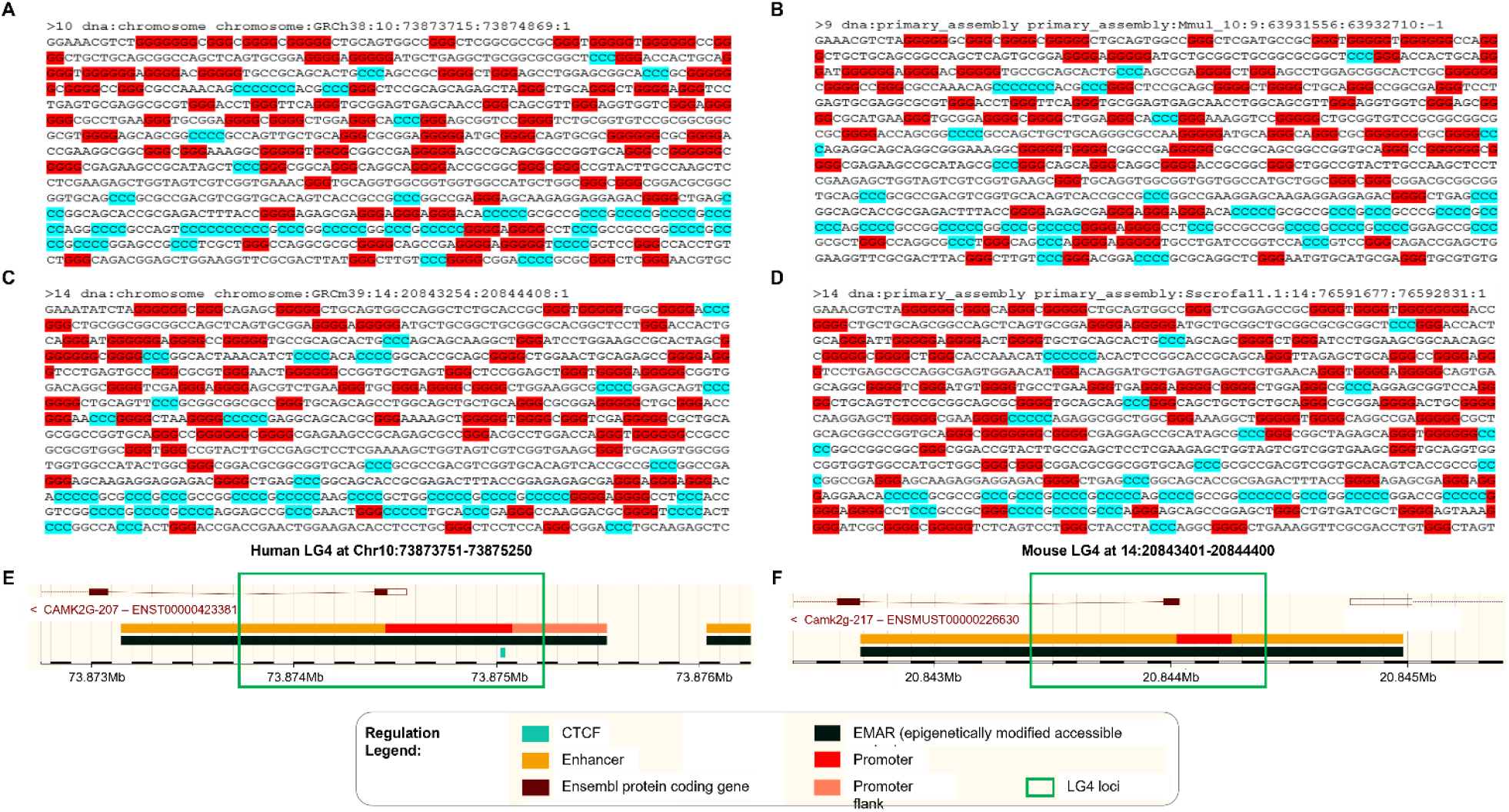
Conservation of the *CAMK2G* LG4 across human, rhesus monkey, mouse, and pig. (**A**) Human LG4 located at GRCh38:10:73873751:73875250. LG4s conserved in (**B**) rhesus monkey (Mmul10:9:63931551:63932050), (**C**) mouse (GRCm39:14:20843401:20844400), and (**D**) pig (Sscrofall.1:14:76591651:76592650) are shown. Sequence shown with 3 or more consecutive Gs highlighted in red and 3 or more consecutive Cs highlighted in blue. Sequence alignments of the human LG4 with each of the conserved LG4s in rhesus monkey, pig, and mouse can be found in **Supplemental Figure 1**. Screenshots of LG4 loci and conserved regulatory elements as depicted in Ensembl^14^: (**E**) Human LG4 located at GRCh38:10:73873751:73875250; (**F**) Mouse LG4 located at (GRCm39:14:20843401:20844400).

### *MAZ* LG4 and its regulatory capacity are conserved across human and mouse

Another conserved LG4 (we deem the *MAZ* LG4) is embedded within the MAZ gene promoter and overlaps an annotated enhancer predicted to regulate over 40 genes, including *MAZ* itself^16^. We find the *MAZ* LG4 neighborhood to be conserved in both human and mouse with both LG4 enhancers predicted to regulate at least seven common genes (**Figure 4**).

**Figure 4.**
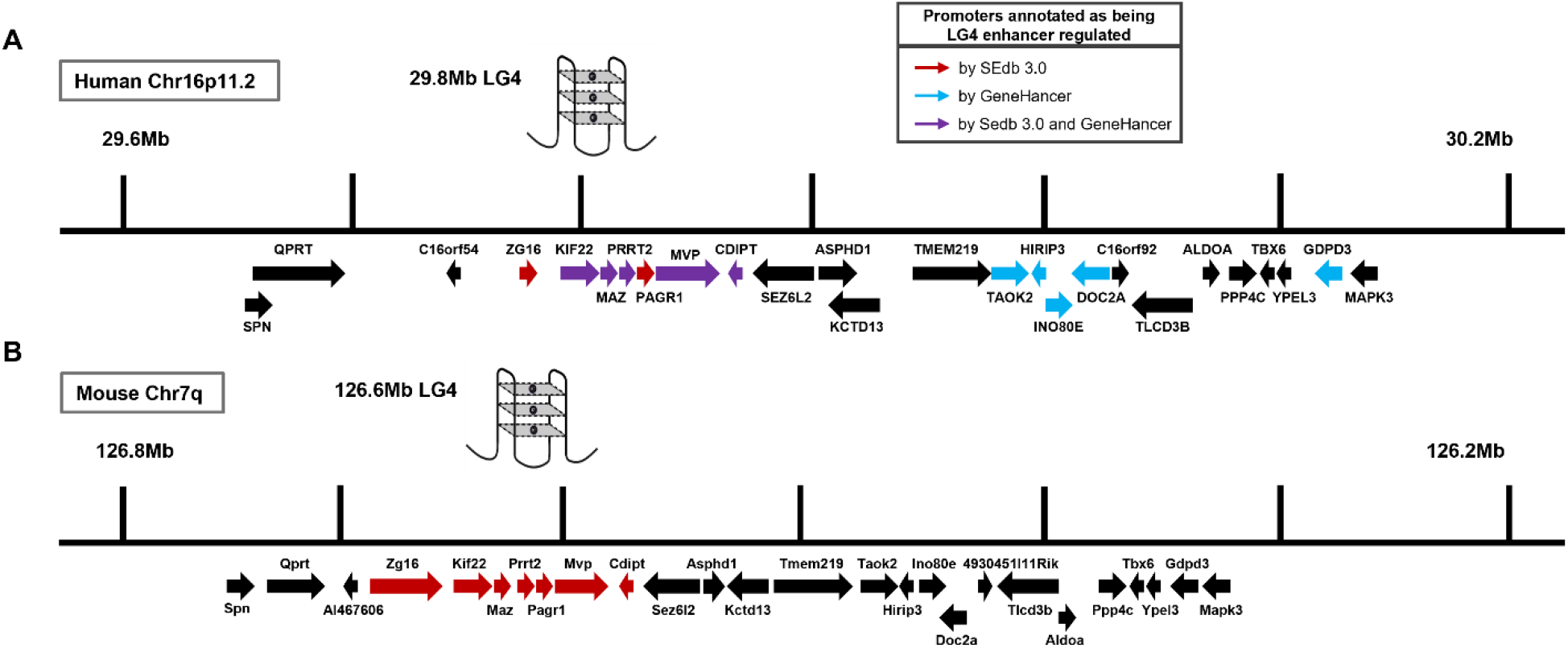
Conservation of the *MAZ* LG4 neighborhood in human and mouse. Genomic neighborhood comparison showing conservation of gene organization surrounding the *MAZ* LG4 Enhancer in the (**A**) human and (**B**) mouse genomes. Conserved gene promoters annotated as being LG4 Enhancer regulated are shown with red, purple, and blue arrows where red = SEdb 3.0^18^ annotated, blue = GeneHancer^16^ annotated, and purple = SEdb 3.0^18^ and GeneHancer^16^ annotated. Black arrows represent conserved genes not annotated as being LG4 Enhancer regulated.

The human *MAZ* LG4 possesses 89% identity to the LG4 found in mouse (**Figure 1B**) and >95% identity to LG4s found in pig and rhesus monkey (**Table 2**). Additionally, this LG4 exhibits consistent positioning relative to the MAZ gene across the aforementioned species, and similarly, the insulated neighborhood surrounding this LG4 is also conserved.

In light of this, we selected the *MAZ* LG4 for further analysis. Specifically, we analyzed the KIF22 (Kinesin Family Member 22) gene promoters from human and mouse to determine their ability to contribute to G4 formation and find them significantly enriched for G triplets (**Figure 5A, B**) suggesting that the *MAZ* LG4 enhancer and *KIF22* promoter may form composite G4s. To test this, we cloned portions of the *MAZ* LG4 in human and the equivalent regions of the *MAZ* LG4 in mouse, as well as their respective *KIF22* promoters, in both the sense and antisense orientations. We then allowed single stranded DNA corresponding to these sequences to fold into composite G4s and successfully demonstrated via electrophoretic mobility shift assay (EMSA) that the *MAZ* LG4 enhancer can directly interact with its respective *KIF22* promoter in both humans and mice (**Figure 5C-E**). We find coincubation of the negative DNA strands of the human *MAZ* LG4 and *KIF22* promoter produces a gel shift indicative of interaction between these sequences exclusively in a G4-permissive environment (+KCl) (**Figure 5D**). Similarly, we find coincubation of the negative DNA strands of the mouse *MAZ* LG4 and *KIF22* promoter produces a potassium-dependent gel shift indicating interaction between these sequences (**Figure 5E**).

**Figure 5.**
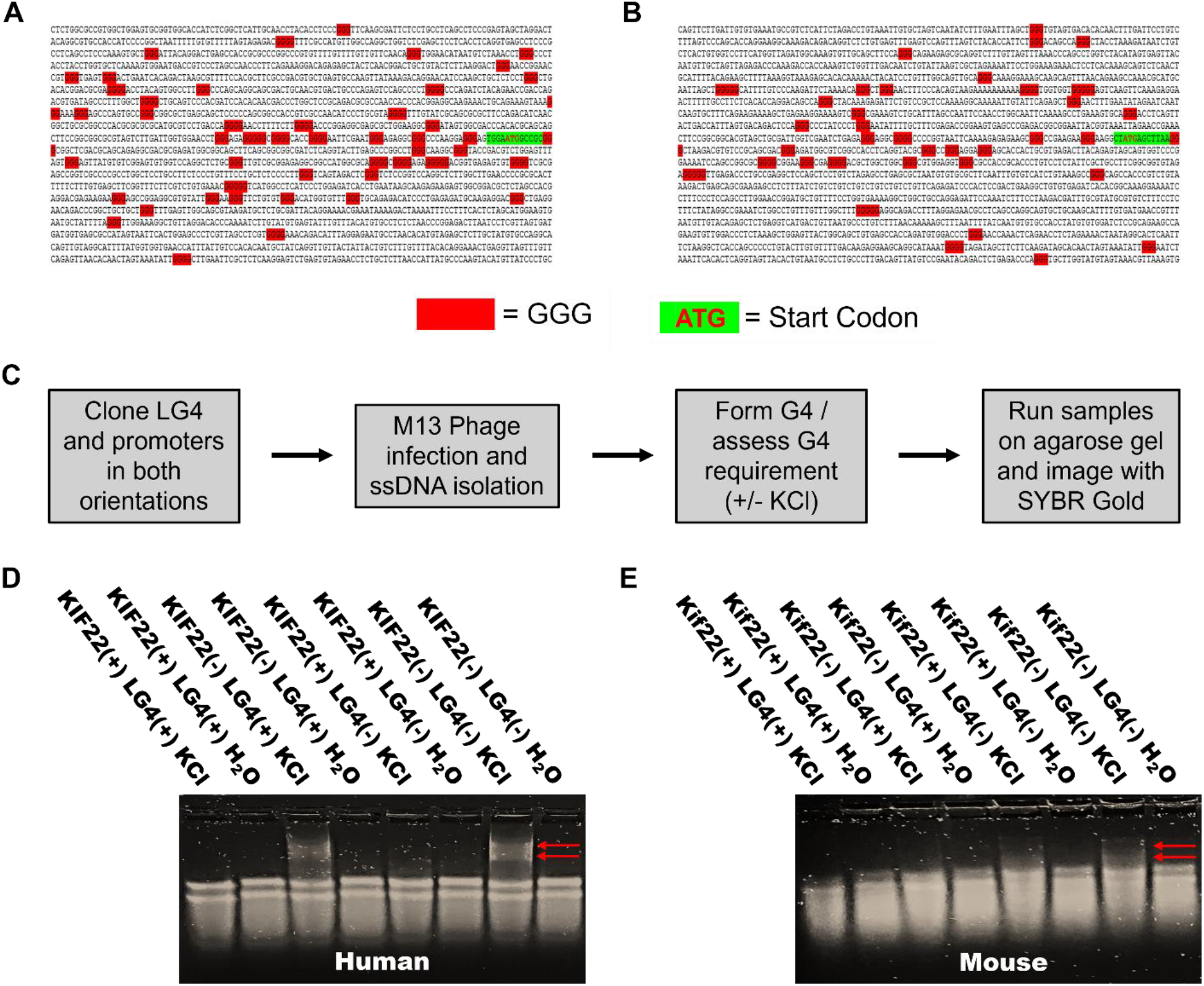
Direct interaction between the *MAZ* LG4 enhancer and *KIF22* promoters in human and mouse. (**A, B**) Both human and mouse *KIF22* promoters are enriched for G triplets (≥3 consecutive genomic guanines and the start codon sequence are highlighted). (**A**) Human *KIF22* promoter located at human Chr16:29789165:29791839:1 (**B**) Mouse *Kif22* promoter Chr7:126640492:126643166:-1. (**C**) Flow chart of EMSA protocol. Adapted from DeMeis et al., 2025. (**D, E**) 1.5% agarose Tris-glycine gel ran at 4°C for 4 hr at 75 V and then stained for 24 hr with SYBR Gold. Constructs were run in either the unfolded (H_2_O) or folded (KCl)[1 M] state as indicated. (**D**) Red arrows denote gel shifts observed when the human *MAZ* LG4 and *KIF22* promoter are folded together. (**E**) Red arrows denote gel shifts observed when the mouse *MAZ* LG4 and *Kif22* promoter are folded together.

## Discussion

In 2020, our lab identified 301 long genomic stretches significantly enriched for minimal G4 motifs which we deemed LG4s. Interestingly, 217 of these LG4s overlap an annotated human enhancer^6^. In 2025, we showed that G4 DNA contained within two of these LG4 enhancers (on human Chr5 and Chr12) directly interact with their target promoters via DNA::DNA interaction^10^.

Since regulatory elements embedded within gene promoters are often evolutionarily conserved^11^, and LG4s are generally associated with human regulatory regions^6^, we hypothesized that LG4s and their regulatory capacity were likely conserved across species. As such, in this study we identified LG4s in the genomes of 13 different species from various taxa (**Table 1**), identified ~2,000 additional LG4s in the human genome (*Homo sapiens*, GRCh38), and found that many of these LG4s are well-conserved in multiple species (**Supplemental Table 4**). In addition, we selected a highly conserved LG4 (**Table 2**) embedded within the MAZ gene promoter (**Figure 1C**) for further analysis. Importantly, not only does the work detailed in this report show that LG4 sequences and their genomic neighborhoods are conserved, but it also demonstrates similar conservation of their regulatory function. Based on a 2015 model of G4-based enhancer:promoter interaction that showed single enhancer:promoter pairs contain minimal, interacting G4 motif components^19^, we analyzed the KIF22 gene promoters in both human and mouse to determine their ability to contribute to G4 formation and found them significantly enriched for G triplets (**Figure 5**). Given that LG4s and their target promoters each contribute a portion of the sequence necessary to form composite G4s^10^, the GGG-rich composition of the *MAZ* LG4 enhancer and *KIF22* promoter suggests that they likely form composite G4s together.

To test this, we cloned an ~500 bp portion of the *MAZ* LG4 in human and the equivalent region of the *MAZ* LG4 in mouse, as well as their respective *KIF22* promoters (in both the sense and antisense orientations). We then generated single-stranded DNAs (ssDNAs) corresponding to each of these sequences and tested their ability to fold into composite G4s in a G4-permissive environment (+KCl). These experiments biochemically confirmed direct interactions between *MAZ* LG4 and *KIF22* promoter DNAs in both humans and mice via EMSA demonstrating functional conservation of this interaction across species. As shown in **Figure 5D** and **E**, EMSA gel shifts indicate interactions between negative strand *MAZ* LG4 and *KIF22* promoter ssDNAs in human and mouse, respectively. Notably, although less pronounced, interactions between negative strand *KIF22* promoter and positive strand LG4 ssDNAs, and conversely, between positive strand *KIF22* promoter and negative strand LG4 ssDNAs were also observed in both human and mouse. That said, observed shifts (1) only occurred in a G4-permissive environment and (2) did not occur when individual *MAZ* LG4 or *KIF22* promoter ssDNAs were allowed to fold in isolation strongly indicating observed shifts do in fact represent composite G4 structures (**Supplemental Figure 8**).

Among the 16 model organism genomes analyzed, we identified LG4s in 13 eukaryotic species including plants, animals, and fungi. Notably we did not identify any LG4s in *S. cerevisiae* or in any prokaryotic genomes, but interestingly, the genome containing the highest percentage of LG4s was that of the unicellular green algae *C. reinhardtii* at 8.25 LG4s per million bp (LG4PM). The only other genomes containing > 1.0 LG4(PM) were *X. tropicalis* (7.24 LG4(PM)), *Z. mays* (4.39 LG4(PM)), and *G. gallus* (2.40 LG4(PM)) suggesting LG4 prevalence is not strongly correlated with any major eukaryotic taxa. That said, all the genomes of multicellular species, excepting *A. thaliana* (0.00 LG4(PM)) and *N. crassa* (0.02 LG4(PM)), were found to contain multiple distinct LG4s (**Table 1**), and generally speaking, we find that less complex multicellular organism genomes contain relatively few LG4s as compared to the genomes of more complex species. Importantly, we find several LG4s associated with species-specific transposable elements^20^ (**data not shown**) and speculate this constitutes the major source of discrepancy between LG4 abundance across species. Further, although the majority of identified LG4s tend to only be conserved in closely related species (also presumably due to LG4s frequently arising from transposons), we identified an LG4 in rhesus monkey (*M. mulatta*) well-conserved in *X. tropicalis* (83.1% identity across 178 bp), as well as two other *X. tropicalis* LG4s conserved in *G. gallus* (91.8% across 60 bp) (**Supplemental Table 4**) suggesting the regulatory functions of these LG4s (and other similarly broadly-conserved LG4s) may be evolutionarily constrained.

Irrespective of our findings, we acknowledge that there are notable limits to this study. In all, we analyzed the genomes of sixteen different species from various taxa, this is a relatively small sample size especially considering the majority of these species are eukaryotes. As such, analyzing the genomes of more species, particularly more prokaryotes, is warranted. Another limit of note is that regulatory elements, and in some cases genes, are not as well-annotated in other organisms as they are in humans leading to the identification of a disproportionate number of overlaps of human LG4s with genes and regulatory elements. Further, the minimally required composition of a functional LG4 remains unclear, and although the version of LG4ID utilized in this work was less stringent than that in our previous study^6^, we cannot discount the possibility of overlooking smaller, less G4-rich LG4s. Finally of note, this study only confirmed conservation of the regulation of one target promoter (*KIF22*) by one LG4 (*MAZ*) in humans and mice, as such further experimentation will clearly be required to ascertain if other LG4 regulations are similarly conserved.

In summary, this work describes LG4s in the genomes of both unicellular and multicellular species including vertebrates, invertebrates, plants, and fungi and shows many of these LG4s (and potentially their regulatory capacities) are highly conserved. Future analyses will likely identify LG4s and LG4 regulations in numerous additional species further supporting the importance of LG4s as hitherto underappreciated regulatory elements.

## Supporting information

Supplemental Table 1. All LG4s identified

Supplemental Table 2. Enhancer Overlaps

Supplemental Table 3. Gene Overlaps

Supplemental Table 4. Unique Alignments

Supplemental Table 5. Oligonucleotide Master List

## Abbreviations

LG4: Long G4-rich region
G4: G-quadruplex
MAZ: Myc-Associated Zinc finger protein
K^+^: Potassium cation
G: Guanine
C: Cytosine
(LG4)PM: LG4# Per Million bp
d/u: data unavailable
CTCF: CCCTC-binding Factor
CAMK2G: Calcium/Calmodulin-Dependent Protein Kinase II Gamma
EMAR: Epigenetically Modified Accessible Region
KIF22: Kinesin Family Member 22
EMSA: Electrophoretic Mobility Shift Assay
KCl: Potassium chloride
BLAST: Basic Local Alignment Search Tool
PCR: Polymerase Chain Reaction
DMSO: Dimethyl sulfoxide

## Declarations

### Ethics approval and consent to participate

Not applicable.

### Consent for publication

Not applicable.

### Availability of data and materials

LG4ID and OverlapID are openly available in the GitHub repositories https://github.com/glen-borchert/LG4ID and https://github.com/glen-borchert/OverlapID, respectively. All corresponding accession links for each species genome, genes, and regulatory features datasets can be found in the **Supplementary Material** file.

### Authors’ contributions

M.H.S. and G.M.B. had full access to all of the data in the study and take responsibility for the integrity of the data and accuracy of the data analysis. This included study concept and design, experimental design and interpretation, data analysis, drafting of manuscript, critical revision of the manuscript for important intellectual content, and general study supervision. J.D.D., C.A.A., M.R.C., T.C.D., K.A.G., G.K.M., J.M.V., M.L.W., S.S.P., and S.Y.A. all contributed directly to the design of specific experiments, data analysis and to overall manuscript editing and approval.

## Acknowledgements

We thank the University of South Alabama College of Medicine Department of Pharmacology for ongoing support.

## References

1. Gellert M, Lipsett MN, Davies DR. Helix formation by guanylic acid. Proc Natl Acad Sci U S A. 1962;48(12):2013–8. doi: 10.1073/pnas.48.12.2013. PubMed PMID: 13947099; PMCID: PMC221115.

2. Kim D, Pertea G, Trapnell C, Pimentel H, Kelley R, Salzberg SL. TopHat2: accurate alignment of transcriptomes in the presence of insertions, deletions and gene fusions. Genome Biol. 2013;14(4):R36. Epub 20130425. doi: 10.1186/gb-2013-14-4-r36. PubMed PMID: 23618408; PMCID: PMC4053844.

3. Sen D, Gilbert W. Formation of parallel four-stranded complexes by guanine-rich motifs in DNA and its implications for meiosis. Nature. 1988;334(6180):364–6. doi: 10.1038/334364a0. PubMed PMID: 3393228.

4. Bochman ML, Paeschke K, Zakian VA. DNA secondary structures: stability and function of G-quadruplex structures. Nat Rev Genet. 2012;13(11):770–80. Epub 20121003. doi: 10.1038/nrg3296. PubMed PMID: 23032257; PMCID: PMC3725559.

5. Kledus F, Dobrovolna M, Mergny JL, Brazda V. Asymmetric distribution of G-quadruplex forming sequences in genomes of retroviruses. Sci Rep. 2025;15(1):76. Epub 20250102. doi: 10.1038/s41598-024-82613-2. PubMed PMID: 39747944; PMCID: PMC11696869.

6. Williams JD, Houserova D, Johnson BR, Dyniewski B, Berroyer A, French H, Barchie AA, Bilbrey DD, Demeis JD, Ghee KR, Hughes AG, Kreitz NW, McInnis CH, Pudner SC, Reeves MN, Stahly AN, Turcu A, Watters BC, Daly GT, Langley RJ, Gillespie MN, Prakash A, Larson ED, Kasukurthi MV, Huang J, Jinks-Robertson S, Borchert GM. Characterization of long G4-rich enhancer-associated genomic regions engaging in a novel loop:loop ‘G4 Kissing’ interaction. Nucleic acids research. 2020;48(11):5907–25. doi: 10.1093/nar/gkaa357.

7. Furlong EEM, Levine M. Developmental enhancers and chromosome topology. Science. 2018;361(6409):1341–5. doi: 10.1126/science.aau0320. PubMed PMID: 30262496; PMCID: PMC6986801.

8. Ptashne M. Gene regulation by proteins acting nearby and at a distance. Nature. 1986;322(6081):697–701. doi: 10.1038/322697a0. PubMed PMID: 3018583.

9. Razin SV, Ulianov SV, Iarovaia OV. Enhancer Function in the 3D Genome. Genes (Basel). 2023;14(6). Epub 20230616. doi: 10.3390/genes14061277. PubMed PMID: 37372457; PMCID: PMC10298269.

10. DeMeis JD, Roberts JT, Delcher HA, Godang NL, Coley AB, Brown CL, Shaw MH, Naaz S, Dahal A, Alqudah SY, Nguyen KN, Nguyen AD, Paudel SS, Shell JE, Patil SS, Dang H, O’Neal WK, Knowles MR, Houserova D, Gillespie MN, Borchert GM. Long G4-rich enhancers target promoters via a G4 DNA-based mechanism. Nucleic Acids Res. 2025;53(2). doi: 10.1093/nar/gkae1180. PubMed PMID: 39658038; PMCID: PMC11754661.

11. Uebbing S, Kocher AA, Baumgartner M, Ji Y, Bai S, Xing X, Nottoli T, Noonan JP. Evolutionary Innovations in Conserved Regulatory Elements Associate With Developmental Genes in Mammals. Mol Biol Evol. 2024;41(10). doi: 10.1093/molbev/msae199. PubMed PMID: 39302728; PMCID: PMC11465374.

12. Perkel J. Democratic databases: science on GitHub. Nature. 2016;538(7623):127–8. doi: 10.1038/538127a. PubMed PMID: 27708327.

13. Haider S, Ballester B, Smedley D, Zhang J, Rice P, Kasprzyk A. BioMart Central Portal--unified access to biological data. Nucleic Acids Res. 2009;37(Web Server issue):W23–7. Epub 20090506. doi: 10.1093/nar/gkp265. PubMed PMID: 19420058; PMCID: PMC2703988.

14. Cunningham F, Amode MR, Barrell D, Beal K, Billis K, Brent S, Carvalho-Silva D, Clapham P, Coates G, Fitzgerald S, Gil L, Girón CG, Gordon L, Hourlier T, Hunt SE, Janacek SH, Johnson N, Juettemann T, Kähäri AK, Keenan S, Martin FJ, Maurel T, McLaren W, Murphy DN, Nag R, Overduin B, Parker A, Patricio M, Perry E, Pignatelli M, Riat HS, Sheppard D, Taylor K, Thormann A, Vullo A, Wilder SP, Zadissa A, Aken BL, Birney E, Harrow J, Kinsella R, Muffato M, Ruffier M, Searle SMJ, Spudich G, Trevanion SJ, Yates A, Zerbino DR, Flicek P. Ensembl 2015. Nucleic acids research. 2015;43(Database issue):D662–9. doi: 10.1093/nar/gku1010.

15. Gao T, He B, Liu S, Zhu H, Tan K, Qian J. EnhancerAtlas: a resource for enhancer annotation and analysis in 105 human cell/tissue types. Bioinformatics. 2016;32(23):3543–51. Epub 20160810. doi: 10.1093/bioinformatics/btw495. PubMed PMID: 27515742; PMCID: PMC5181530.

16. Fishilevich S, Nudel R, Rappaport N, Hadar R, Plaschkes I, Iny Stein T, Rosen N, Kohn A, Twik M, Safran M, Lancet D, Cohen D. GeneHancer: genome-wide integration of enhancers and target genes in GeneCards. Database: the journal of biological databases and curation. 2017;2017. doi: 10.1093/database/bax028.

17. Altschul SF, Gish W, Miller W, Myers EW, Lipman DJ. Basic local alignment search tool. Journal of molecular biology. 1990;215(3):403–10. doi: 10.1016/S0022-2836(05)80360-2.

18. Song S, Liu L, Feng C, Xie L, Zhang G, Zhang Y, Gao Y, Yin M, Tang X, Pei W, Song C, Liu R, Li C. SEdb 3.0: a comprehensive super-enhancer database across multiple species. Nucleic Acids Res. 2025. Epub 20251208. doi: 10.1093/nar/gkaf1294. PubMed PMID: 41359035.

19. Hegyi H. Enhancer-promoter interaction facilitated by transiently forming G-quadruplexes. Scientific reports. 2015;5:9165–. doi: 10.1038/srep09165.

20. Jurka J. Repbase update: a database and an electronic journal of repetitive elements. Trends Genet. 2000;16(9):418–20. doi: 10.1016/s0168-9525(00)02093-x. PubMed PMID: 10973072.

