## Supplemental Figures for "Conservation of Long G4-rich (LG4) genomic enhancer regulations"

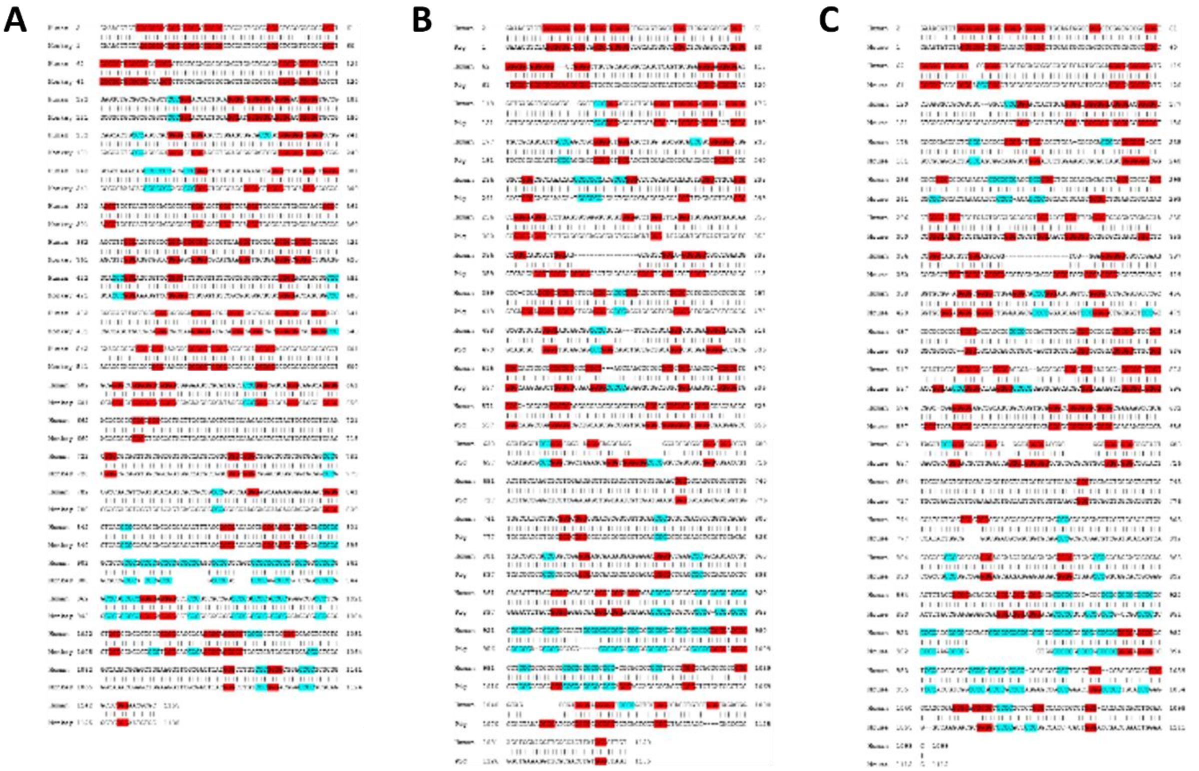

**Supplemental Figure 1. Sequence alignments of the human *CAMK2G* LG4 against the same conserved LG4s in rhesus monkey, pig, and mouse.** Sequences are shown with 3 or more consecutive Guanines highlighted in red and 3 or more consecutive Cytosines highlighted in blue. BLAST LG4 sequence alignments of human LG4 (GRCh38:10:73873751:73875250) are depicted as follows: (**A**) human LG4 vs. LG4 conserved in rhesus monkey (Mmul_10:9:63931551:63932050), (**B**) human LG4 vs. LG4 conserved in pig (Sscrofall.1:14:76591651:76592650), and (**C**) human LG4 vs. LG4 conserved in mouse (GRCm39:14:20843401:20844400).

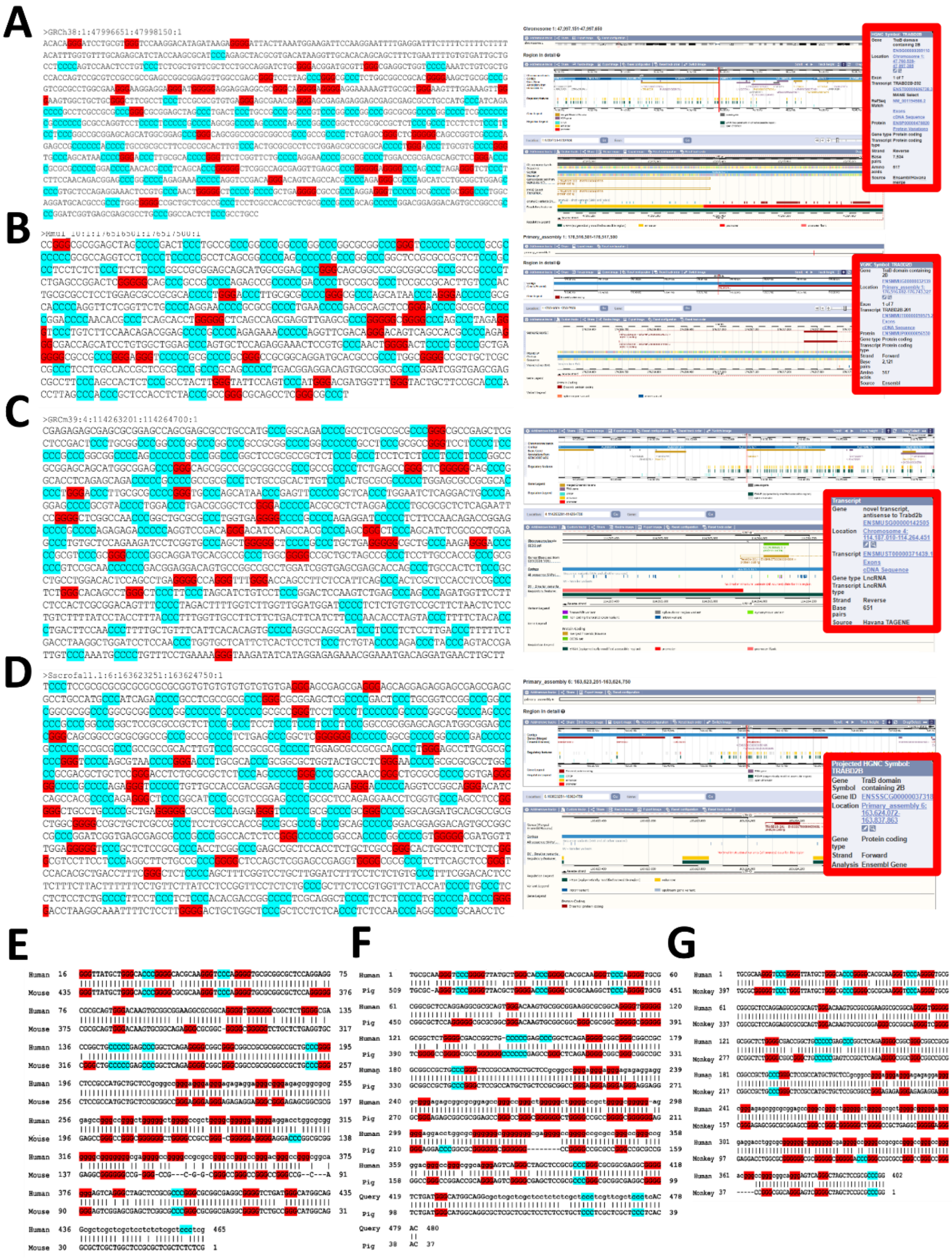


**Supplemental Figure 2. Conservation of the *TRABD2B* LG4 and associated regulatory features across human, rhesus monkey, mouse, and pig.** (**A**) Human LG4 located at GRCh38:1:47997151:47997650. Sequence is shown (left) with 3 or more consecutive Guanines highlighted in red and 3 or more consecutive Cytosines highlighted in blue. Screenshots of LG4 loci as depicted in Ensembl[14] are also included with overlapping genes and enhancers indicated (right). Corresponding conserved loci in (**B**) rhesus monkey (Mmul_10:1:176516501:176517500), (**C**) mouse (GRCm39:4:114263201:114264700), and (**D**) pig (Sscrofall.1:6:163623251:163624750) are shown. BLAST sequence alignments with 3 or more Gs and Cs highlighted as in **A** of (**E**) human and mouse loci, (**F**) human and pig loci, and (**G**) human and monkey loci have also been included.

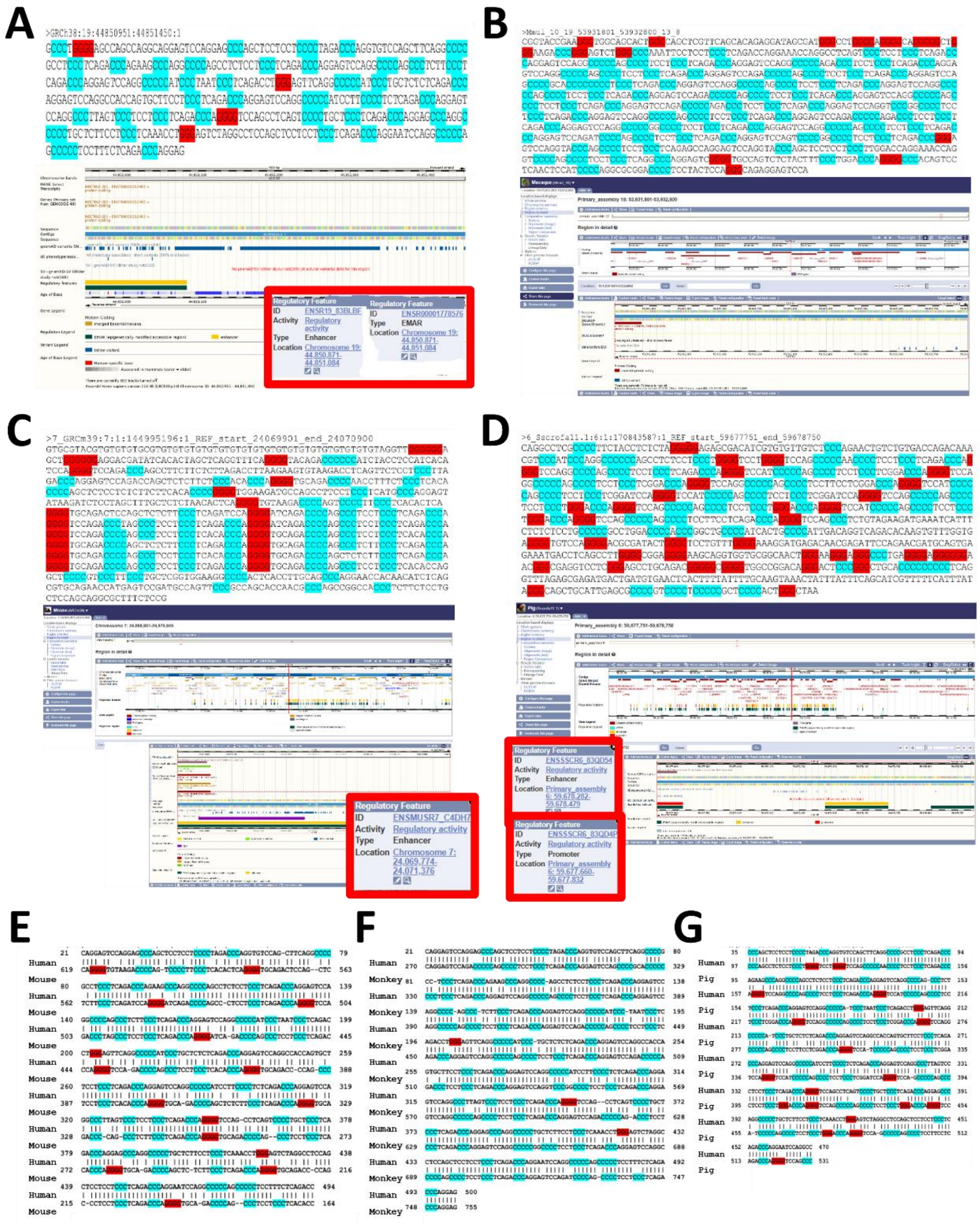


**Supplemental Figure 3. Conservation of the *NECTIN2* LG4 and associated regulatory features across human, rhesus monkey, mouse, and pig.** (**A**) Human LG4 located at GRCh38:19:44850951:44851450. Sequence is shown (top) with 3 or more consecutive Guanines highlighted in red and 3 or more consecutive Cytosines highlighted in blue. Screenshots of LG4 loci as depicted in Ensembl[14] are also included with overlapping genes and enhancers indicated (bottom). Corresponding conserved loci in (**B**) rhesus monkey (Mmul_10:19:53931801:53932800), (**C**) mouse (GRCm39:7:24069901:24070900), and (**D**) pig (Sscrofall.1:6:59677751:59678750) are shown. BLAST sequence alignments with 3 or more Gs and Cs highlighted as in **A** of (**E**) human and mouse loci, (**F**) human and monkey loci, and (**G**) human and pig loci have also been included.

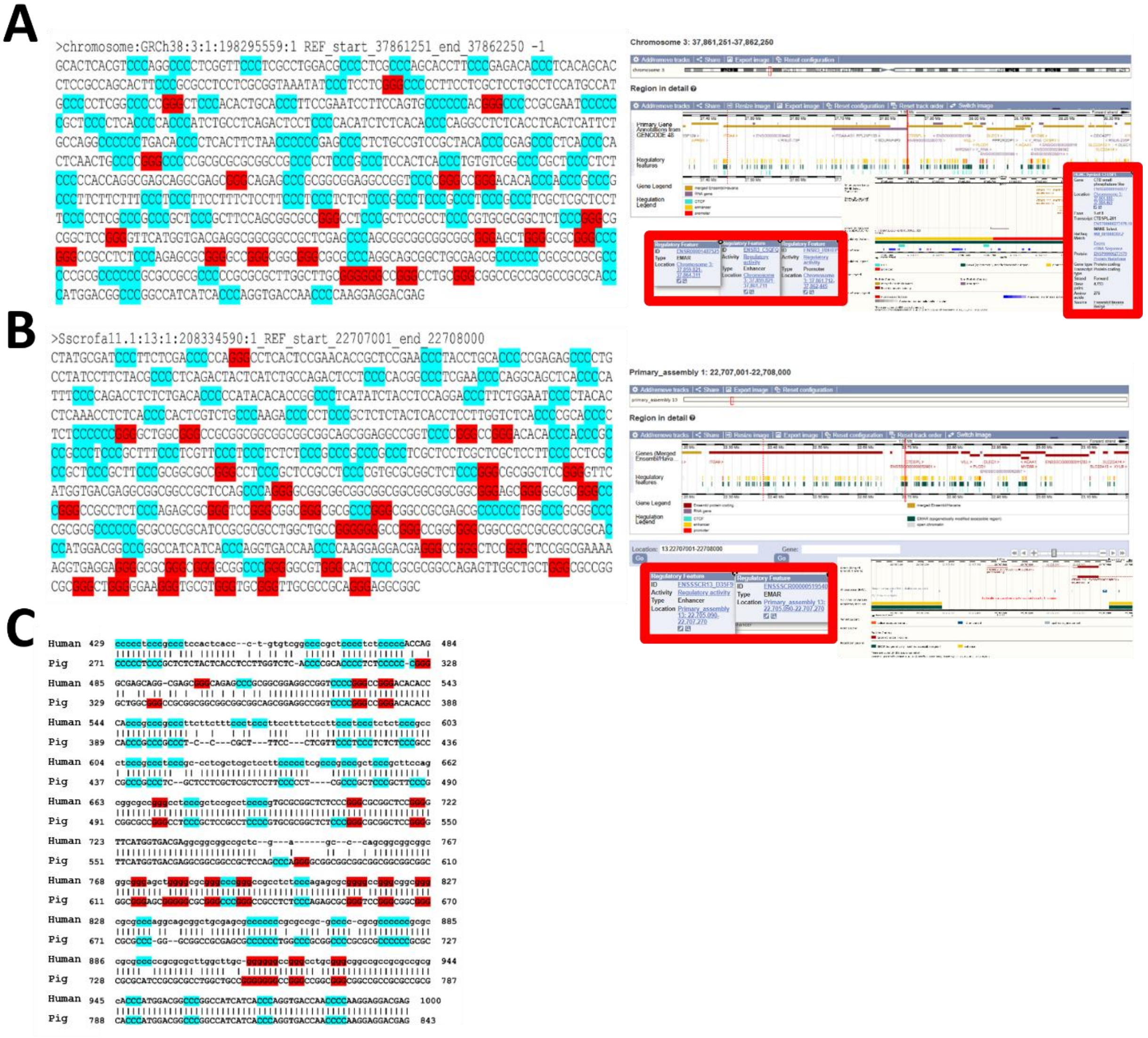


**Supplemental Figure 4. Conservation of the *CTDSPL* LG4 and associated regulatory features across human and pig.** (**A**) Human LG4 located at h38:3:37861251:37862250. Sequence is shown (left) with 3 or more consecutive Guanines highlighted in red and 3 or more consecutive Cytosines highlighted in blue. Screenshots of LG4 loci as depicted in Ensembl[14] are also included with overlapping genes and enhancers indicated (right). Corresponding conserved loci in (**B**) pig (Sscrofall.1:13:22707001:22708000) are shown. BLAST sequence alignments with 3 or more Gs and Cs highlighted as in A of (**C**) human and pig loci have also been included.

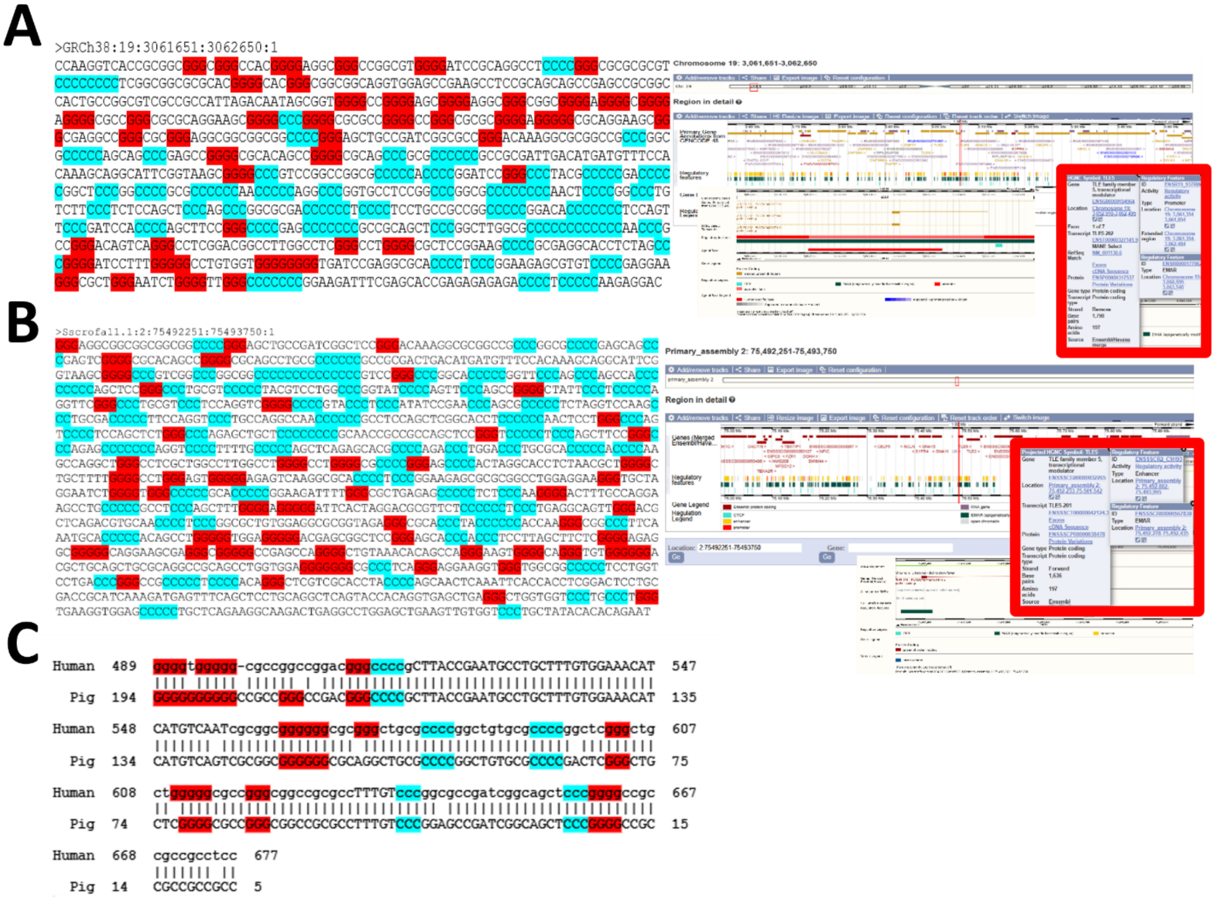


**Supplemental Figure 5. Conservation of the *TLE5* LG4 and associated regulatory features across human and pig.** (**A**) Human LG4 located at GRCh38:19:3061651:3062650. Sequence is shown (left) with 3 or more consecutive Guanines highlighted in red and 3 or more consecutive Cytosines highlighted in blue. Screenshots of LG4 loci as depicted in Ensembl[14] are also included with overlapping genes and enhancers indicated (bottom). Corresponding conserved loci in (**B**) pig (Sscrofall.1:19:3061651:3062650) are shown. BLAST sequence alignments with 3 or more Gs and Cs highlighted as in **A** of (**C**) human and pig loci have also been included.

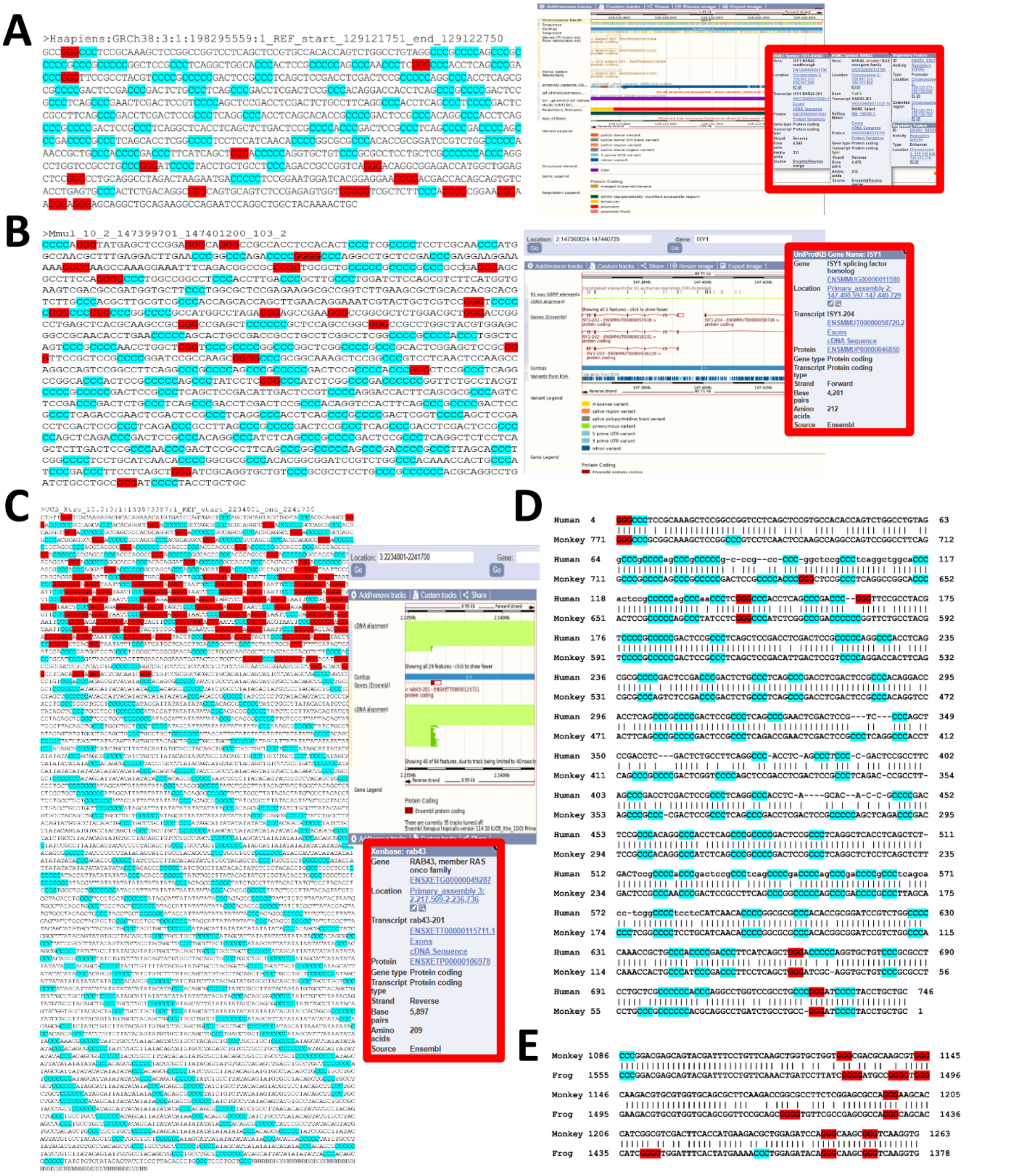


**Supplemental Figure 6. Conservation of the *RAB43* LG4 and associated regulatory features across human, rhesus monkey, and frog.** (**A**) Human LG4 located at GRCh38:3:129121751:129122750. Sequence is shown (left) with 3 or more consecutive Guanines highlighted in red and 3 or more consecutive Cytosines highlighted in blue. Screenshots of LG4 loci as depicted in Ensembl[14] are also included with overlapping genes and enhancers indicated (right). Corresponding conserved loci in (**B**) rhesus monkey (Mmul_10:10:147399701:147401200), (**C**) frog (UCB_Xtro_10.0:3:2234801:2241700) are shown. BLAST sequence alignments with 3 or more Gs and Cs highlighted as in **A** of (**D**) human and monkey loci, and (**E**) monkey and frog loci have also been included.

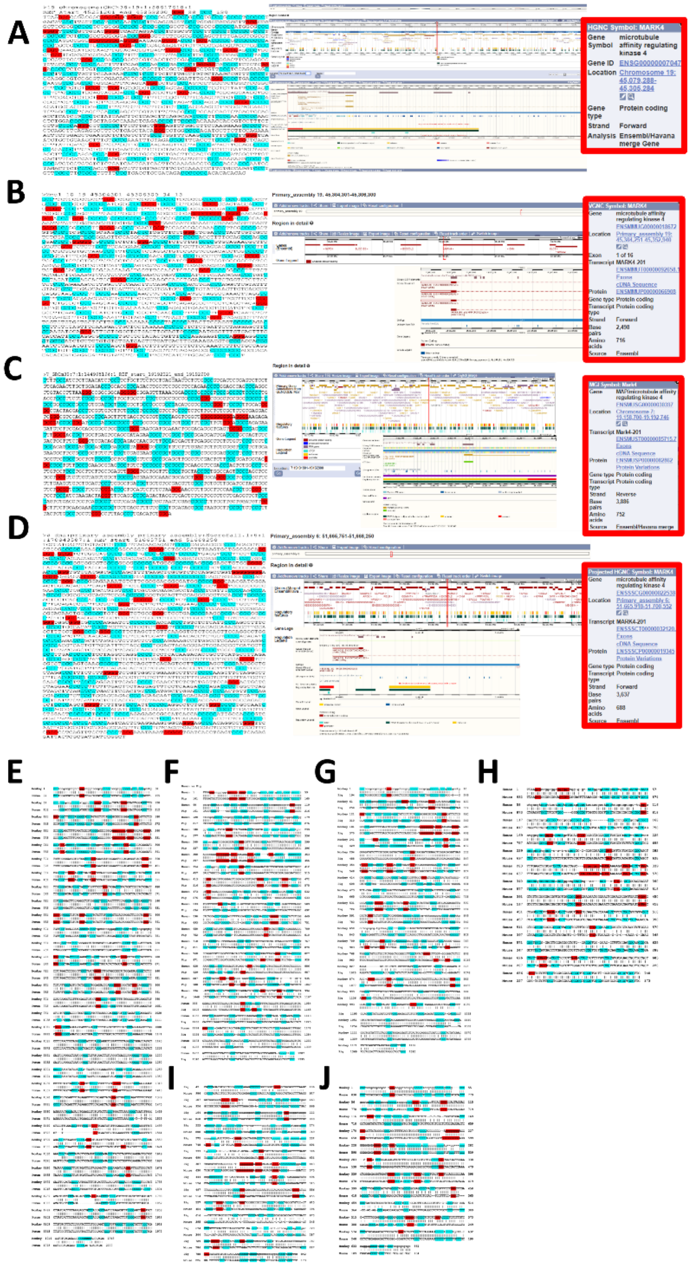


**Supplemental Figure 7. Conservation of the *MARK4* LG4 and associated regulatory features across human, rhesus monkey, mouse, and pig.** (**A**) Human LG4 located at GRCh38:19:45251201:45253200. Sequence is shown (left) with 3 or more consecutive Guanines highlighted in red and 3 or more consecutive Cytosines highlighted in blue. Screenshots of LG4 loci as depicted in Ensembl[14] are also included with overlapping genes and enhancers indicated (right). Corresponding conserved loci in (**B**) rhesus monkey (Mmul_10:19:45304301:45306300), (**C**) mouse (GRCm39:7:19191901:19192900), and (**D**) pig (Sscrofa11.1:6:51665751:51668250) are shown. BLAST sequence alignments with 3 or more Gs and Cs highlighted as in **A** of (**E**) human and monkey loci, (**F**) human and pig loci, (**G**) monkey and pig loci, (**H**) human and mouse loci, (**I**) pig and mouse loci, and (**J**) monkey and mouse loci have also been included.

A

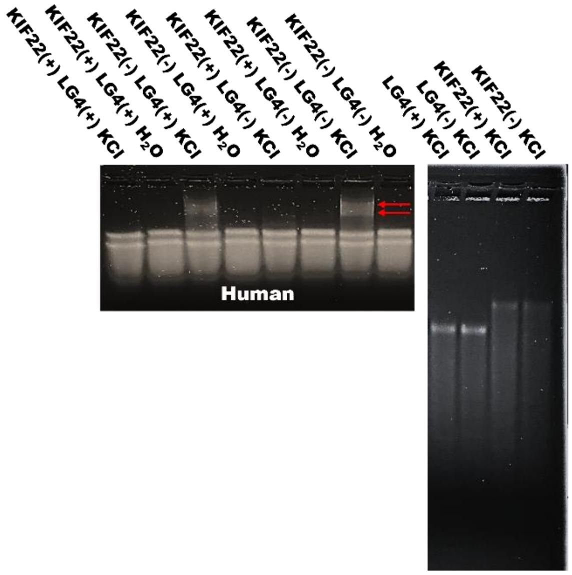


B

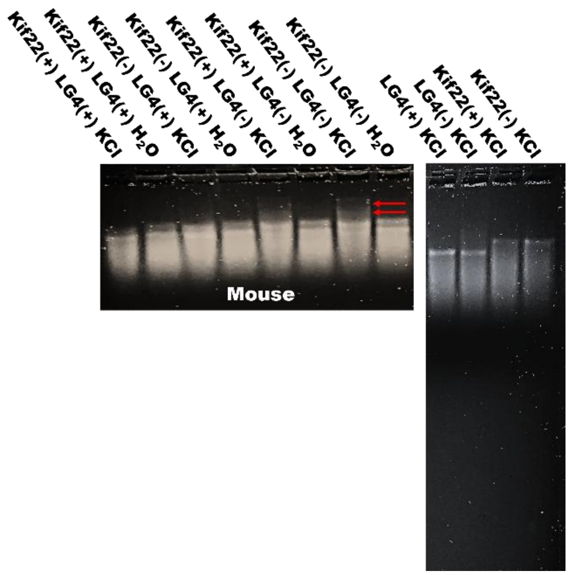


**Supplemental Figure 8. Direct interaction between the *MAZ* LG4 enhancer and *KIF22* promoters in human and mouse.** 1.5% agarose Tris-glycine gels ran at 4°C for 4 hr at 75 V and then stained for 24 hr with SYBR Gold. Constructs were run in either the unfolded ($H_2O$) or folded (KCl)[1 M] state as indicated. Identically treated *KIF22* or LG4 ssDNAs run individually are shown to the right of gels containing these same ssDNAs run together following coincubation. (**A**) Red arrows denote gel shifts observed when the human *MAZ* LG4 and *KIF22* promoter are folded together. (**B**) Red arrows denote gel shifts observed when the mouse *MAZ* LG4 and *Kif22* promoter are folded together.
