## Supplementary Material for "Conservation of Long G4-rich (LG4) genomic enhancer regulations"

The genomes used in this study were from Ensembl Release 113, Ensembl Plants Release 59, Ensembl Metazoa Release 59, Ensembl Fungi Release 59, and Ensembl Bacteria Release 59. The gene sets for each species were obtained from Ensembl BioMart corresponding to the same release as each of the respective genomes. The regulatory features were obtained via: Ensembl BioMart (*Homo sapiens* and *Mus musculus*), GeneHancer (*Homo sapiens*), and EnhancerAtlas (*Gallus gallus*, *Sus scrofa*, *Danio rerio*, *Drosophila melanogaster*, and *Caenorhabditis elegans*).

***Xenopus tropicalis*:**

<https://ftp.ensembl.org/pub/release-113/fasta/xenopus_tropicalis/dna/Xenopus_tropicalis.UCB_Xtro_10.0.dna.toplevel.fa.gz>

Genes:

<http://sep2024.archive.ensembl.org/biomart/martview/64d9b91383ef926277ad2fde87d78487?VIRTUALSCHEMANAME=default&ATTRIBUTES=xtropicalis_gene_ensembl.default.feature_page.ensembl_gene_id|xtropicalis_gene_ensembl.default.feature_page.ensembl_gene_id_version|xtropicalis_gene_ensembl.default.feature_page.ensembl_transcript_id|xtropicalis_gene_ensembl.default.feature_page.ensembl_transcript_id_version&FILTERS=&VISIBLEPANEL=mainpanel>

***Zea mays*:**

<https://ftp.ebi.ac.uk/ensemblgenomes/pub/release-59/plants/fasta/zea_mays/dna/Zea_mays.Zm-B73-REFERENCE-NAM-5.0.dna.toplevel.fa.gz>

Genes:

<http://may2024-plants.ensembl.org/biomart/martview?VIRTUALSCHEMANAME=plants_mart&ATTRIBUTES=zmays_eg_gene.default.feature_page.ensembl_gene_id|zmays_eg_gene.default.feature_page.ensembl_transcript_id&FILTERS=&VISIBLEPANEL=mainpanel>

***Macaca mulatta*:**

<https://ftp.ensembl.org/pub/release-113/fasta/macaca_mulatta/dna/Macaca_mulatta.Mmul_10.dna.toplevel.fa.gz>

Genes:

<http://oct2024.archive.ensembl.org/biomart/martview/bdc25ff98883669a3b42028fb9e45c4d?VIRTUALSCHEMANAME=default&ATTRIBUTES=mmulatta_gene_ensembl.default.feature_page.ensembl_gene_id|mmulatta_gene_ensembl.default.feature_page.ensembl_gene_id_version|mmulatta_gene_ensembl.default.feature_page.ensembl_transcript_id|mmulatta_gene_ensembl.default.feature_page.ensembl_transcript_id_version&FILTERS=&VISIBLEPANEL=mainpanel>

***Gallus gallus*:**

<https://ftp.ensembl.org/pub/release-113/fasta/gallus_gallus/dna/Gallus_gallus.bGalGal1.mat.broiler.GRCg7b.dna.toplevel.fa.gz>

Genes:

<http://oct2024.archive.ensembl.org/biomart/martview/bdc25ff98883669a3b42028fb9e45c4d?VIRTUALSCHEMANAME=default&ATTRIBUTES=ggallus_gene_ensembl.default.feature_page.ensembl_gene_id|ggallus_gene_ensembl.default.feature_page.ensembl_gene_id_version|ggallus_gene_ensembl.default.feature_page.ensembl_transcript_id|ggallus_gene_ensembl.default.feature_page.ensembl_transcript_id_version&FILTERS=&VISIBLEPANEL=mainpanel>

Regulatory Features:

<http://enhanceratlas.org/data/download/species_enh_bed.tar.gz>

***Homo sapiens*:**

<https://ftp.ensembl.org/pub/release-113/fasta/homo_sapiens/dna/Homo_sapiens.GRCh38.dna.primary_assembly.fa.gz>

Genes:

<http://oct2024.archive.ensembl.org/biomart/martview/bdc25ff98883669a3b42028fb9e45c4d?VIRTUALSCHEMANAME=default&ATTRIBUTES=hsapiens_gene_ensembl.default.feature_page.ensembl_gene_id|hsapiens_gene_ensembl.default.feature_page.ensembl_gene_id_version|hsapiens_gene_ensembl.default.feature_page.ensembl_transcript_id|hsapiens_gene_ensembl.default.feature_page.ensembl_transcript_id_version&FILTERS=&VISIBLEPANEL=mainpanel>

Regulatory Features (Ensembl BioMart):

<http://oct2024.archive.ensembl.org/biomart/martview/4215b0dae59ecd91b85bd3925ba5c60e?VIRTUALSCHEMANAME=default&ATTRIBUTES=hsapiens_regulatory_feature.default.regulatory_feature.chromosome_name|hsapiens_regulatory_feature.default.regulatory_feature.chromosome_start|hsapiens_regulatory_feature.default.regulatory_feature.chromosome_end|hsapiens_regulatory_feature.default.regulatory_feature.feature_type_name&FILTERS=&VISIBLEPANEL=mainpanel>

Regulatory Features (GeneHancer):

<https://www.genecards.org/Guide/DatasetRequest>

***Mus musculus*:**

<https://ftp.ensembl.org/pub/release-113/fasta/mus_musculus/dna/Mus_musculus.GRCm39.dna.primary_assembly.fa.gz>

Genes:

<http://oct2024.archive.ensembl.org/biomart/martview/bdc25ff98883669a3b42028fb9e45c4d?VIRTUALSCHEMANAME=default&ATTRIBUTES=mmusculus_gene_ensembl.default.feature_page.ensembl_gene_id|mmusculus_gene_ensembl.default.feature_page.ensembl_gene_id_version|mmusculus_gene_ensembl.default.feature_page.ensembl_transcript_id|mmusculus_gene_ensembl.default.feature_page.ensembl_transcript_id_version&FILTERS=&VISIBLEPANEL=mainpanel>

Regulatory Features:

<http://oct2024.archive.ensembl.org/biomart/martview/4215b0dae59ecd91b85bd3925ba5c60e?VIRTUALSCHEMANAME=default&ATTRIBUTES=mmusculus_regulatory_feature.default.regulatory_feature.chromosome_name|mmusculus_regulatory_feature.default.regulatory_feature.chromosome_start|mmusculus_regulatory_feature.default.regulatory_feature.chromosome_end|mmusculus_regulatory_feature.default.regulatory_feature.feature_type_name&FILTERS=&VISIBLEPANEL=mainpanel>

***Chlamydomonas reinhardtii*:**

<https://ftp.ebi.ac.uk/ensemblgenomes/pub/release-59/plants/fasta/chlamydomonas_reinhardtii/dna/Chlamydomonas_reinhardtii.Chlamydomonas_reinhardtii_v5.5.dna.toplevel.fa.gz>

Genes:

<http://may2024-plants.ensembl.org/biomart/martview?VIRTUALSCHEMANAME=plants_mart&ATTRIBUTES=creinhardtii_eg_gene.default.feature_page.ensembl_gene_id|creinhardtii_eg_gene.default.feature_page.ensembl_transcript_id&FILTERS=&VISIBLEPANEL=mainpanel>

***Sus scrofa*:**

<https://ftp.ensembl.org/pub/release-113/fasta/sus_scrofa/dna/Sus_scrofa.Sscrofa11.1.dna.toplevel.fa.gz>

Genes:

<http://sep2024.archive.ensembl.org/biomart/martview/64d9b91383ef926277ad2fde87d78487?VIRTUALSCHEMANAME=default&ATTRIBUTES=sscrofa_gene_ensembl.default.feature_page.ensembl_gene_id|sscrofa_gene_ensembl.default.feature_page.ensembl_gene_id_version|sscrofa_gene_ensembl.default.feature_page.ensembl_transcript_id|sscrofa_gene_ensembl.default.feature_page.ensembl_transcript_id_version&FILTERS=&VISIBLEPANEL=mainpanel>

Regulatory Features:

<http://enhanceratlas.org/data/download/species_enh_bed.tar.gz>

***Danio rerio*:**

<https://ftp.ensembl.org/pub/release-113/fasta/danio_rerio/dna/Danio_rerio.GRCz11.dna.primary_assembly.fa.gz>

Genes:

<http://oct2024.archive.ensembl.org/biomart/martview/bdc25ff98883669a3b42028fb9e45c4d?VIRTUALSCHEMANAME=default&ATTRIBUTES=drerio_gene_ensembl.default.feature_page.ensembl_gene_id|drerio_gene_ensembl.default.feature_page.ensembl_gene_id_version|drerio_gene_ensembl.default.feature_page.ensembl_transcript_id|drerio_gene_ensembl.default.feature_page.ensembl_transcript_id_version&FILTERS=&VISIBLEPANEL=mainpanel>

Regulatory Features:

<http://enhanceratlas.org/data/download/species_enh_bed.tar.gz>

***Drosophila melanogaster*:**

<https://ftp.ensembl.org/pub/release-113/fasta/drosophila_melanogaster/dna/Drosophila_melanogaster.BDGP6.46.dna.toplevel.fa.gz>

Genes:

<http://may2024-metazoa.ensembl.org/biomart/martview?VIRTUALSCHEMANAME=metazoa_mart&ATTRIBUTES=dmelanogaster_eg_gene.default.feature_page.ensembl_gene_id|dmelanogaster_eg_gene.default.feature_page.ensembl_transcript_id&FILTERS=&VISIBLEPANEL=mainpanel>

Regulatory Features:

<http://enhanceratlas.org/data/download/species_enh_bed.tar.gz>

***Caenorhabditis elegans*:**

<https://ftp.ensembl.org/pub/release-113/fasta/caenorhabditis_elegans/dna/Caenorhabditis_elegans.WBcel235.dna.toplevel.fa.gz>

Genes:

<http://oct2024.archive.ensembl.org/biomart/martview/bdc25ff98883669a3b42028fb9e45c4d?VIRTUALSCHEMANAME=default&ATTRIBUTES=celegans_gene_ensembl.default.feature_page.ensembl_gene_id|celegans_gene_ensembl.default.feature_page.ensembl_transcript_id&FILTERS=&VISIBLEPANEL=mainpanel>

Regulatory Features:

<http://enhanceratlas.org/data/download/species_enh_bed.tar.gz>

***Schizophyllum commune*:**

<https://ftp.ebi.ac.uk/ensemblgenomes/pub/release-59/fungi/fasta/fungi_basidiomycota1_collection/schizophyllum_commune_h4_8_gca_000143185/dna/Schizophyllum_commune_h4_8_gca_000143185.v1.0.dna.toplevel.fa.gz>

Genes:

<https://ftp.ebi.ac.uk/ensemblgenomes/pub/release-62/fungi/gtf/schizophyllum_commune/Schizophyllum_commune.GCA000143185v2.62.gtf.gz>

***Neurospora crassa*:**

<https://ftp.ebi.ac.uk/ensemblgenomes/pub/release-59/fungi/fasta/neurospora_crassa/dna/Neurospora_crassa.NC12.dna.toplevel.fa.gz>

Genes:

<http://may2024-fungi.ensembl.org/biomart/martview?VIRTUALSCHEMANAME=fungi_mart&ATTRIBUTES=ncrassa_eg_gene.default.feature_page.ensembl_gene_id|ncrassa_eg_gene.default.feature_page.ensembl_transcript_id&FILTERS=&VISIBLEPANEL=mainpanel>

***Arabidopsis thaliana*:**

<https://ftp.ebi.ac.uk/ensemblgenomes/pub/release-59/plants/fasta/arabidopsis_thaliana/dna/Arabidopsis_thaliana.TAIR10.dna.toplevel.fa.gz>

***Haloferax volcanii*:**

<https://ftp.ensemblgenomes.ebi.ac.uk/pub/release-59/bacteria//fasta/bacteria_0_collection/haloferax_volcanii_ds2_gca_000025685/dna/Haloferax_volcanii_ds2_gca_000025685.ASM2568v1.dna.toplevel.fa.gz>

***Saccharomyces cerevisiae*:**

<https://ftp.ebi.ac.uk/ensemblgenomes/pub/release-59/fungi/fasta/saccharomyces_cerevisiae/dna/Saccharomyces_cerevisiae.R64-1-1.dna.toplevel.fa.gz>
